# The impact of resource provisioning on the epidemiological responses of different parasites

**DOI:** 10.1101/2021.05.22.445253

**Authors:** Diana Erazo, Amy B. Pedersen, Andy Fenton

## Abstract

1. Events such as anthropogenic activities and periodic tree masting can alter resource provisioning in the environment, directly affecting animals, and potentially impacting the spread of infectious diseases in wildlife. The effect of these supplemental resources on infectious diseases can manifest through different pathways, affecting host susceptibility, transmission and host demography.
2. To date however, empirical research has tended to examine these different pathways in isolation, for example by quantifying the effects of provisioning on host behaviour in the wild or changes in immune responses in controlled laboratory studies. Further, while theory has investigated the interactions between these pathways, thus far this work has focussed on a narrow subset of pathogen types, typically directly-transmitted microparasites. Given the diverse ways that provisioning can affect host susceptibility, contact patterns or host demography, we may expect the epidemiological consequences of provisioning to depend on key aspects of parasite life-history, such as the duration of infection and transmission mode.
3. We developed a suite of generic epidemiological models to compare how resource provisioning alters responses for different parasites that vary in their biology (micro- and macro-parasite), transmission mode (direct, environmental, and vector transmitted) and duration of infection (acute, latent, and chronic). Next, we parameterised these different parasite types using data from the diverse parasite community of wild wood mice as a case study.
4. We show there are common epidemiological responses to host resource provisioning across all parasite types examined. In particular, the response to provisioning could be driven in opposite directions, depending on which host pathways (contact rate, susceptibility or host demography) are most altered by the addition of resources to the environment. Broadly, these responses were qualitatively consistent across all parasite types, emphasising the importance of identifying general trade-offs between provisioning-altered parameters.
5. Despite the qualitative consistency in responses to provisioning across parasite types, we found notable quantitative differences between parasites, suggesting specific epidemiological outcomes could strongly depend on parasite type, infection duration and permanency of recovery, and whether the parasite is directly, environmentally, or vector transmitted. These analyses therefore highlight the importance of knowing key specific aspects of host-parasite biology, such as host contact behaviours, parasite interactions with the host immune system, and how resource availability shapes host demographics, in order to understand and predict epidemiological responses to provisioning for any specific host-parasite system.

## Introduction

The availability of food resources is a key environmental factor that impacts the demography and behaviour of animals, but also has the ability to impact host-parasite interactions (Bradley & Altizer, 2007; Robb, McDonald, Chamberlain, & Bearhop, 2008). Natural events, such as periodic tree masting, and human activities such as agricultural expansion or the purposeful feeding of wildlife, can alter host-parasite interactions through several distinct pathways, acting across multiple ecological scales (Jensen, 1982; Oro, Genovart, Tavecchia, Fowler, & Martínez-Abraín, 2013). At the individual scale, food supplementation (or ‘provisioning’) can improve host body condition and immune responses, which can result in lower susceptibility to infection, shorter-lasting infections, and better health outcomes for the host (Becker, Streicker, & Altizer, 2015; Cypher & Frost, 1999). For example, a study on lace monitors foraging on urban waste suggested that body condition improved as a result of provisioning, and this improvement was associated with a lower intensity of blood-borne parasites compared to unprovisioned individuals (Jessop, Smissen, Scheelings, & Dempster, 2012). At the between-host scale, hosts can aggregate around resources, thereby increasing contact rates and transmission (Boutin, 1990), as seen by the aggregation of European greenfinches around supplemental bird feeders, which increased the transmission of *Trichomonas gallinae* (Lawson et al., 2012). Finally, at the host population scale, parasite prevalence and transmission potential (as measured by the basic reproductive number; R_0_) can increase through host demographic responses to provisioning, through the addition of new susceptible individuals to the population via new births or immigration, which can potentially increase the likelihood of parasite persistence (Boutin, 1990; Krebs et al., 1995; Nagy & Holmes, 2005). Therefore, resource provisioning can affect parasite dynamics through separate pathways involving changes in host susceptibility, behaviour and/or demography. Together, these changes can lead to either net increases or decreases in infection risk, and impact the epidemiological outcomes in different directions, depending on the balance of how each pathway is impacted by provisioning.

Given the many ways in which provisioning can affect infectious disease epidemiology, mechanistic models can be invaluable tools to explore how various provisioning-altered parameters relating to host demography, contact behaviour, and host immune defence interact to affect the dynamics of parasite infection and spread. In particular, previous theoretical studies demonstrated that a range of parasite responses (i.e. changes in prevalence and basic reproductive number, R_0_) could arise from coupling functional responses of host birth and death, host susceptibility and contact rate parameters to the level of resource provisioning (Becker & Hall, 2014). For instance, increases in both host density and contact rates due to resource provisioning boost R_0_ (McCallum, Barlow, & Hone, 2001), when host immunity is not directly affected. However, if a host with access to improved nutritional resources was better able to protect itself from infection due a strengthened immune response, then the parasite R_0_ is significantly reduced. Additionally, parasite extinction could even occur at intermediate provisioning levels, with invasion (R_0_) and persistence only occurring at low or high provisioning levels (Becker et al., 2015). Thus, by showing that parasite persistence changes along a gradient of resource provisioning, depending on the underlying effects of resources on the host. Therefore, modelling approaches are valuable tools for understanding the opposing patterns in disease outcomes observed in nature.

To date, however, theory has focussed on a narrow subset of parasite types, typically directly-transmitted microparasites that fit the classic ‘Susceptible-Infected’ (SI) or ‘Susceptible-Infected-Recovered’ (SIR) frameworks. Given the diverse ways that provisioning can affect proximate factors relating host physiology/immunology, contact patterns or host demography, we may expect the ultimate epidemiological consequences of provisioning to depend on key aspects of parasite life-history, such as the duration of infection, parasite type and transmission mode. For instance, environmentally transmitted or vector-borne parasites could exhibit different responses to provisioning-induced changes in host population size and contact structure, compared to directly-transmitted parasites. Furthermore, how macroparasites, rather than microparasites, are impacted by provisioning of the host remains poorly understood, as impacts on burden-dependent processes (Anderson & May, 1978; Dobson & Hudson, 1992) or altered environments for parasite reproduction and survival (Seppälä, Liljeroos, Karvonen, & Jokela, 2008) may introduce additional responses not seen for microparasites. To understand how different parasite types may vary in their overall responses to host resource provisioning, we constructed a series of five epidemiological models, which represent a range of parasites that vary in their biology (micro- and macroparasite), transmission mode (direct, environmental, and vector transmitted) and duration of infection (acute, latent, and chronic). Next, we parameterised these different parasite types using data from the diverse parasite community of wild wood mice (*Apodemus sylvaticus*) as a case study. We demonstrate that there are some generalities in how different parasites respond to resource provisioning of the host that are consistent across parasite types and transmission modes. However, we also found notable differences in the magnitude of these responses by the different parasites. These analyses therefore highlight the importance of identifying specific aspects of host and parasite biology, to properly infer the effects of provisioning for any specific host-parasite system.

## Methods

### Model construction

We constructed five compartmental models, each describing a different, but common parasite transmission structure: (i) macroparasite, (ii) microparasite SIR (susceptible-infected-recovered), (iii) environmentally transmitted microparasite SIS (susceptible-infected), (iv) microparasite SAL (susceptible-active-latent), and (v) vector-borne microparasite SIS (Figure 1). The specific details of each model are given in the Supplementary Material, but they all followed standard structures well-known from the epidemiological literature (e.g., (Anderson & May, 1978, 1981, 1991) Importantly, all five models contained parameters that incorporated the effects of increased resource provisioning of the host on the three key, separate pathways: (1) host demography (through long-term changes to the host’s carrying capacity, *K*), (2) contact behaviour (*α*), and (3) immune defence (through changes to host susceptibility, *δ*). In particular we assumed the host’s carrying capacity (*K*) and contact rate (*α*) would increase with resource provision to the host, while host susceptibility (*δ*) would decrease as hosts had more access to supplemental nutrition (as in Becker & Hall (2014) and Becker, Streicker & Altizer (2015)).

**Figure 1.**
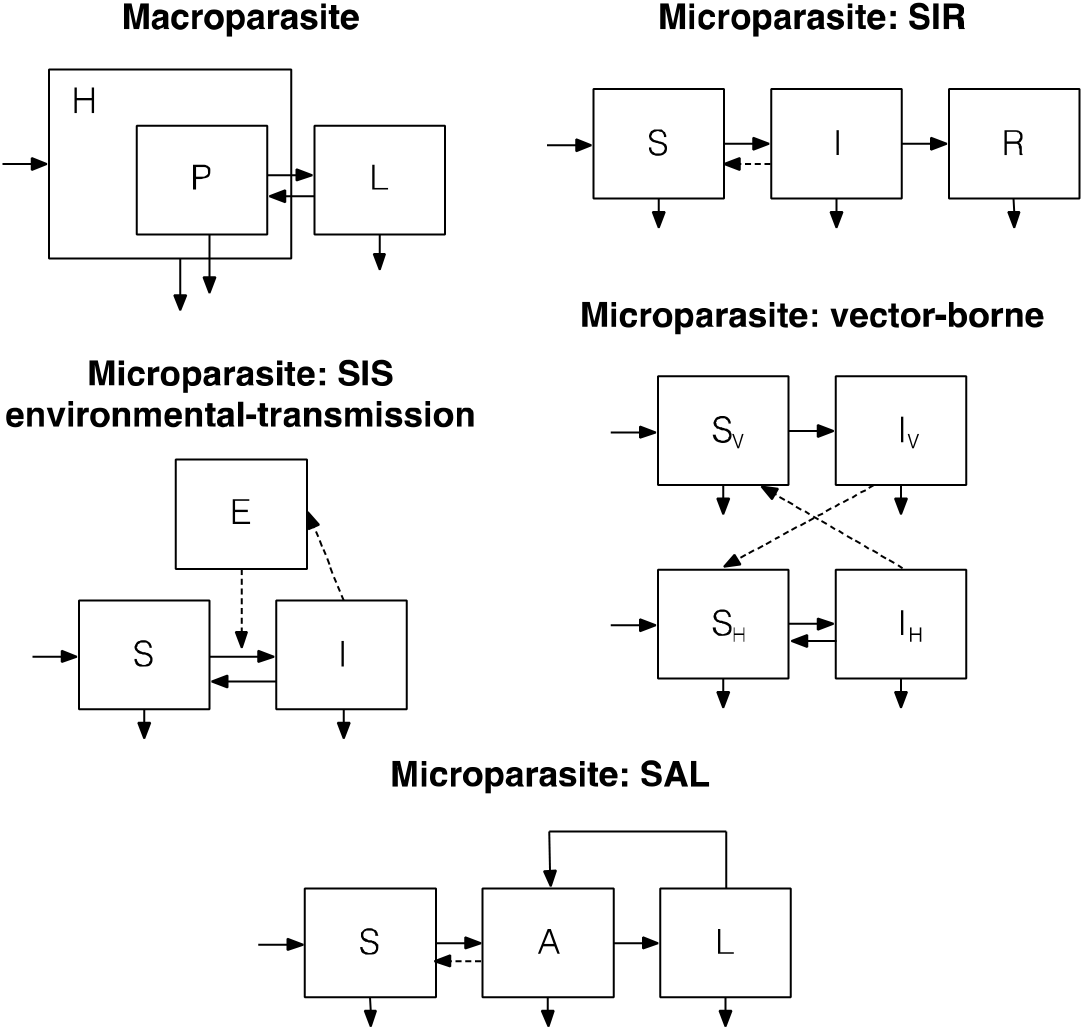
Schematic diagram of the macroparasite and microparasite models. Boxes represent populations in each system. For microparasite models, hosts were classified by their infection status, therefore *S, I, R, A* and *L* represented susceptible, infected, recovered, acute and latent populations, respectively. For the vector-borne parasite, the subscripts V and H denote classes relating to the vector and host populations respectively. For the macroparasite model, only one host population (*H*) was considered, *P* represents the size of the adult parasite population and *L* represents the size of the environmental pool of infective parasite larvae. Arrows represent populations transitions and dashed arrows represent the mechanism through susceptible population gets infected.

To capture the effects of resource provisioning on both host contact rate and susceptibility, we followed the approach developed by Becker & Hall (2014), where provisioning is represented by the parameter *ρ*. Here *ρ* = 0 represents no supplemental resource provisioning (hence, all parameters are at baseline values), while increasing values of *ρ* correspond to increasing levels of resource provisioning, for example due to human-provided food sources, land-use change increasing access to novel resources, or periodic access to resources, such as through tree masting. As in Becker & Hall (2014), parameter responses to provisioning were assumed to have a monotonic and saturating behaviour that depends on the parameter’s sensitivity to provisioning, represented by *θ_x_*, allowing parameters to scale from linear forms (*θ_x_* = 0) to a quickly saturating shape (high values of *θ_x_*). Since the contact rate between hosts (*α*) is expected to increase with provisioning (Becker et al., 2015; Boutin, 1990), we used the following functional form:

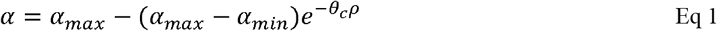

Because susceptibility to infection (*δ*) is hypothesised to decrease with provisioning (Becker et al., 2015; Sweeny et al., 2019) we used the following function:

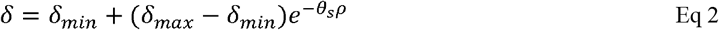

where *α_min_* and *δ_min_* are the minimum values, and *α_max_* and *δ_max_* are the maximum values that contact rate and susceptibility could take. Thus, if *ρ*= 0, *δ* = *δ_max_* and *α* = *α_min_*, the parameter baseline values. In an intensive resource provisioned scenario (high *ρ*) the maximum contact rate was assumed to be twice the baseline value (*α_max_* = 2*α_min/bl_*) and the minimum susceptibility was assumed to be half the baseline value 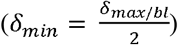.

We investigated the net effect of host resource provisioning for each of the above parasite types as the difference in either mean parasite burdens for macroparasites or parasite prevalence for microparasites at equilibrium, between arbitrarily intensive provisioning (*ρ* = 1) and no provisioning (*ρ* = 0) scenarios. Here a value greater than 0 for these differences implies a net increase in overall prevalence or mean parasite burden due to provisioning, whereas a value less than 0 implies a net reduction due to provisioning. We explored how these changes for each parasite depended on the balance of the host defence and host contact sensitivities to resource provisioning (i.e., within *θ_s_* – *θ_c_* parameter space). To simplify the analysis and presentation, we incorporated the effects of provisioning on host demography by varying host carrying capacity from *K* to 3*K*, to reflect long-term increases in overall food availability due to increased provisioning (Becker et al., 2015; Ozoga & Verme, 1982). Hence, this analysis allowed us to explore how the overall changes due to provisioning vary with the sensitivity of the individual host demographic, contact rate or susceptibility parameters to provisioning.

### Model parameterization

While our intention is to explore generic responses of each parasite type to provisioning, we fit each model to field data, to ensure we were using biologically plausible values. Specifically we parameterised each model using a specific parasite species from our four-year longitudinal dataset of 1453 individually-tagged wood mice (*Apodemus sylvaticus*) captured across four 70×70m grids, at Haddon Wood on the Wirral, UK between June 2009 and December 2012 (Knowles, Fenton, & Pedersen, 2012; Sweeny, Albery, Venkatesan, Fenton, & Pedersen, 2021). Wood mice in these populations are commonly infected by five different species of parasites, each of which corresponded to the different modelling frameworks described above (see Table 1 for details): *Heligmosomoides polygyrus* (macroparasite)*, Eimeria hungaryensis* (SIS), Wood Mouse Herpes Virus (WMHV - SAL)*, Bartonella* spp. (vector-borne), *Trypanosoma grosi* (vector-borne), and Cowpox virus (SIR). Full details of model parametrisation are provided in the Supplementary Material but, briefly, parameters were estimated using adaptive Monte Carlo Markov Chain Metropolis-Hastings (MCMC-MH) using the R package fitR (Camacho & Funk, 2019), assuming uniform priors. Model fitting was carried out in two stages. First, host demographic parameters were estimated by fitting the predicted total number of mice per week to the observed number of mice captured per week. Next, we estimated parasite infection-related parameters for the various parasite transmission models (see Table 1), through fitting each model individually. For the microparasite models, we used data on seroprevalence for Wood Mouse Herpes Virus (Knowles et al., 2012) and PCR diagnostics for *Trypanosoma grosi* and *Bartonella* spp. (Knowles et al., 2013; Withenshaw, Devevey, Pedersen, & Fenton, 2016), while we used faecal egg or oocyst counts (measured as eggs or oocysts per gram of faeces) for the gastrointestinal nematode *H. polygyrus* and the coccidian *E. hungaryensis*, respectively (Knowles et al., 2013). Because the seroprevalence of Cowpox was very low in the wood mouse population (<5%), the infection-related parameters were estimated using data from previous literature (Telfer et al., 2002); see Supplementary Material.

**Table 1.** Parasites from the *A. sylvaticus* system: model, type, transmission mode, diagnostic, and parameter values. K = 42.12* in all parasite models. *Parameter baseline values. See Supplementary Material for details of parameter estimation.

| Parasite<br>(Model) | Type | Transmission<br>mode | Diagnostic | $\alpha_{min}$<br>$\rho = 0^*$ | $\alpha_{max}$<br>$\rho = 1$ | $\delta_{min}$<br>$\rho = 1$ | $\delta_{max}$<br>$\rho = 0^*$ |
| --- | --- | --- | --- | --- | --- | --- | --- |
| **Cowpox virus**<br>(SIR) | Virus | Contact | Seroprevalence | 0.1 | 0.2 | 0.053 | 0.105 |
| ***Eimeria***<br>***hungaryensis***<br>(SIS) | Coccidian | Environmental | Faecal eggs<br>count | 0.00001 | 0.0002 | 0.048 | 0.096 |
| **Herpesvirus**<br>(SAL) | Virus | Contact | Seroprevalence | 0.1 | 0.2 | Male to male:<br>0.128<br>Female to male:<br>0.212<br>Male to female:<br>0.074<br>Female to female:<br>0.078 | Male to male:<br>0.255<br>Female to male:<br>0.424<br>Male to female:<br>0.148<br>Female to<br>female:<br>0.156 |
| ***Bartonella* spp.**<br>(vector-borne) | Bacteria | Vector | PCR | 0.01 | 0.02 | 0.408 | 0.815 |
| ***Trypanosoma***<br>***grosi***<br>(vector-borne) | Protozoan | Vector | PCR | 0.01 | 0.02 | 0.053 | 0.106 |
| ***Heligmosomoides***<br>***polygyrus***<br>(macroparasite) | Nematode | Environmental | Faecal eggs<br>count | 0.001 | 0.002 | 0.103 | 0.205 |

Estimated parameter values in Table 1 were assumed to be the baseline values (δ = *δ_max_* and α = *α_min_*) corresponding to the scenario of no supplemental resource provisioning (*ρ*= 0). Under the scenario of arbitrarily intensive resource provisioning (*ρ* = 1), parameter values for *δ* and *α* were calculated using Equations 1 and 2 (Becker & Hall, 2014).

## Results

For all 5 parasite species, we found the same qualitative responses of parasite infection (either prevalence or mean burden) to host resource provisioning; all parasite infection measures increased when contact rate was highly sensitive to provisioning (*θ_c_* → 5) and susceptibility sensitivity to provisioning was minimal (*θ_s_* = 0) (green regions, Figure 2). Hence, as may be expected, rapid increases in host contact rate (and hence transmission) due to provisioning (high *θ_c_*), coupled with a negligible response of host immunity to provisioning (low *θ_s_*) result in high levels of infection for all parasite types. Conversely, prevalence and infection burden for all parasites was the most reduced in the opposite scenario, when host susceptibility was highly sensitive to provisioning and the impact of provisioning on contact rate was minimal (*θ_c_* = 0, *θ_s_* → 5; red regions, Figure 2). Hence, rapid increases in immunity due to provisioning (high *θ_s_*), coupled with negligible changes to contact rate (low *θ_c_*), result in low levels of infection, and this was true across all parasite types.

**Figure 2.**
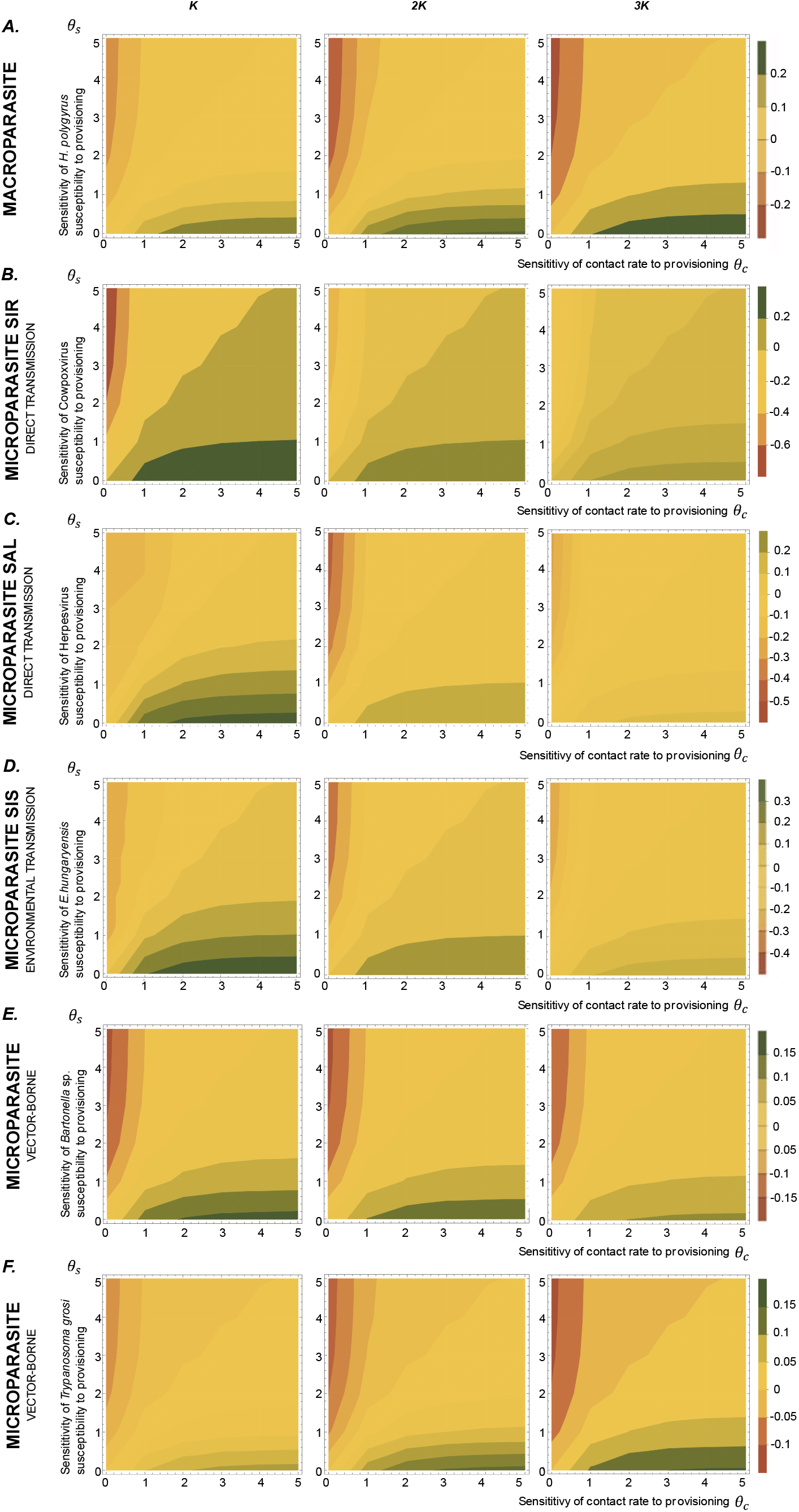
Resource-provision impacts parasite infection and burdens. The A row shows macroparasite mean burden change and B row to F row represent microparasites’ prevalence changes (the difference in equilibrium prevalence between provisioned (*ρ*= 1) and un-provisioned scenarios (*ρ*= 0)). Columns illustrate different carrying capacities, varying from *K* to 3*K*, representing increasing levels of host demographic response to provisioning. For all figures, *X-*axis represents the sensitivity of contact rate to provisioning (*θ_c_*) and *y-*axis is the sensitivity of host susceptibility to provisioning (*θ_s_*).

However, although all parasites showed the same qualitative responses as *θ_s_* and *θ_c_* varied, the different parasite types differed considerably in their quantitative responses to provisioning (i.e., in the magnitude of changes in prevalence or mean infection burden, as indicated by the intensities of colours in Figure S1, which presents the same results as Figure 2, but re-expressed so the changes in prevalences are plotted on the same scale for all microparasites). For clarity, we initially describe these results assuming no host demographic change to provisioning (carrying capacity fixed at the baseline value of *K*; Fig S1 left-hand column), but below we consider the effects of incorporating longer-term host demographic changes due to resource provisioning. When the carrying capacity is fixed at *K*, the directly-transmitted SIR microparasite showed the strongest responses overall to provisioning, with ~20% increase in prevalence due to provisioning for high *θ_c_* (rapid increase in contact rate due to provisioning) and low *θ_s_* (negligible change in immunity due to provisioning), and ~70% decrease in prevalence due to provisioning for low *θ_c_* and high *θ_s_* (Figure S1, row *B*). The SAL and SIS microparasites also showed a reasonably strong (~10%) increase in prevalence due to provisioning for high *θ_c_* and low *θ_s_*, but very little change in prevalence for all other combinations of parameters (Figure S1, rows *C, D*). Comparatively however, the two vector-borne parasites presented very minor quantitative variations in prevalence due to provisioning across all combinations of *θ_c_* and *θ_s_* (Figure S1, rows *E, F*).

When we included the effect of longer-term host demographic responses to provisioning, modelled as fixed changes in host carrying capacity, the varying parasite types responded differently. As carrying capacity was increased from *K* to 3*K*, changes in macroparasite mean burden became more extreme (Figure 2, row *A*); with both increases in burden under high contact rate sensitivity (*θ_c_* → 5), and decreases in burden under high susceptibility sensitivity (*θ_s_* → 5), becoming intensified with higher host carrying capacities. Conversely, the opposite pattern occurred for the microparasite SIR model, where the change in prevalence became less dramatic as carrying capacity increased (e.g., whereas high *θ_c_* was associated with >20% increase in prevalence when the carrying capacity was kept at *K* (Figure 2 row *B*, left-hand plot), increasing the carrying capacity to 3*K* resulted in <10% increase in prevalence (Figure 2 row *B*, right-hand plot)). The SAL and SIS models however showed mixed responses to rises in carrying capacity; the magnitude of positive changes in prevalence due to provisioning (i.e., in the region of high *θ_c_* and low *θ_s_*) were neutralized as carrying capacity increased for both parasites (Figure 2 rows *C* and *D*), whereas the region of negative changes in prevalence due to provisioning (i.e., in the region of low *θ_c_* and high *θ_s_*) initially intensified as carrying capacity was increased from *K* to 2*K*, but then diminished as it was increased further to 3*K*.

Finally, the two vector-borne parasites showed different predicted responses to changes in host demography; *Bartonella* spp. (relatively high baseline prevalence, reflecting high estimated baseline host susceptibility; see Supplementary Material) was predicted to show broadly the same pattern regardless of carrying capacity variations, with both positive and negative changes in prevalence being approximately equal as carrying capacity changed (Figure 2, row *E*). On the other hand, changes in host prevalence for *T. grosi* (low baseline prevalence, with low estimated baseline host susceptibility) became intensified as carrying capacity increased, similarly to that seen for changes in macroparasite mean burden (Figure 2, row *F*).

## Discussion

Overall, our analyses revealed common epidemiological responses of a wide range of different parasite types to resource provisioning of their host. In particular, for all parasites, we demonstrated that responses to provisioning could be driven in opposite directions dependent on which host pathways are most altered by provisioning (Becker & Hall, 2014): increases in contact rate and population size due to provisioning could boost parasite prevalence, whereas decreases in host susceptibility might decrease it. However, we also found some notable differences, particularly in the magnitude of responses by the different parasites, suggesting specific epidemiological outcomes could strongly depend on parasite type (micro- or macroparasite), the duration of infection (whether SIS, SIR or SAL), and whether the parasite is directly, environmentally, or vector-borne transmitted (Altizer et al., 2018). These analyses therefore highlight the importance of understanding the key aspects of host-parasite biology, to properly infer effects of resource provisioning on host-parasite interactions.

We found that different parasite types, such as the macroparasite *H. polygyrus* and vector-borne parasites, which strongly depend on host demographic factors such as changes in host population size, were likely to show extreme changes as carrying capacity increases due to provisioning. Our analyses demonstrated that the macroparasite, and the vector-borne parasite with low baseline prevalence, showed the greatest changes in mean worm burden and prevalence, respectively, to host demographic responses to provisioning. In general, for directly transmitted parasites, however, increases in host carrying capacity due to provisioning translated into limited increases, and sometimes even net decreases, in infection prevalence. These predictions are supported by observations from larger vampire bat (*Desmodus rotundus*) colonies in livestock areas, where provisioning was found to impact host population size, but no effect of rabies virus exposure was observed (Streicker et al., 2012). However, our analysis also showed that directly transmitted parasites that do not confer lasting immunity (i.e., the SIS model) presented the lowest parasite prevalence at intermediate population sizes, particularly if host susceptibility rapidly declines with provisioning (high *θ_s_*). This finding supports previous research suggesting that at an intermediate demographic response to provisioning, parasite persistence (as measured by its R_0_) attains its minimum, although susceptibility would have to be more affected by provisioning compared contact rate (Becker et al., 2015).

Prevalence was predicted to increase for both directly and environmentally-transmitted microparasites, when the response of host aggregation behaviour to provisioning is stronger compared to host immune defence and demographic responses. An example of this scenario has been reported in European greenfinches (*Chloris chloris*), in which aggregation around bird feeders increases transmission of the parasite *Trichonomas gallinae* (Lawson et al., 2012). Similarly in Elk (*Cervus elaphus*) populations, supplemental feed stations were shown to increase contact rates and lead to higher *Brucella abortis* exposure (Cross, Edwards, Scurlock, Maichak, & Rogerson, 2007). In contrast, host susceptibility can also be more affected by improved nutrition compared to changes in contact rate. One study on lace monitors (*Varanus varius*) foraging on human garbage, found that body condition improvement was associated with lower intensity of blood parasites compared to non-provisioned individuals (Jessop et al., 2012). Another example of this scenario occurs in wood mice infected with *H. polygyrus*, in which resource provisioned mice were less susceptible to infection, cleared worms more efficiently, and maintained better body condition (Sweeny et al., 2019). These examples suggest that increased host aggregation, and therefore increased transmission, is not always an outcome of provisioning, or at least not sufficient to overcome potential beneficial impacts of provisioning on the host’s physiology. Indeed, changes in host contact rates due to resource provisioning are likely to depend explicitly on the spatial arrangement of resources, thus, when food is aggregated, contact rate between hosts could increase, as observed in urban feral cats showing smaller foraging ranges and higher territory overlap compared to rural cats (Schmidt, Lopez, & Collier, 2007). However, if food is more evenly scattered across the landscape, contact rate could actually decrease with provisioning. Spatially explicit models could be useful for studying how resource provisioning modifies contact rates and therefore parasite persistence (Plowright et al., 2011).

In this study, we assumed that any impact of resource provisioning on host body condition would be positive, and therefore decrease host susceptibility to parasite infection. However, this may not always be the case, and it is important to consider both resource quantity and quality; it is quite possible that although resource quantity might increase, for example through increased availability of human garbage, quality could be poorer compared to a baseline scenario, thereby increasing host susceptibility to infection (Maggini, Wintergerst, Beveridge, & Hornig, 2007; Van Heugten, Coffey, & Spears, 1996). For instance, rock iguanas feeding from carbohydrate-rich food sources provided by tourists, were associated with changes in nutritional status and higher hookworm burdens compared to non-supplemented individuals (Knapp et al., 2013). Furthermore, although we considered a range of parasite types, we did not consider either multi-host systems or parasites with complex life cycles involving intermediate hosts. Such systems would require consideration of how those alternative host species themselves respond to resource provisioning, potentially resulting in more complex net responses of such parasites to provisioning. Future theoretical studies on the role of provisioning on parasite transmission more generally could therefore involve even more complex systems than those considered here.

In conclusion, we demonstrate that there are general responses to provisioning that are consistent across a wide range of parasite types and transmission modes; thus, our work contributes to the understanding and comparison of increased provisioning impacts on different parasites in the same host species to allow comparison. In particular, we demonstrated that considering general trade-offs of provisioning-altered parameters is crucial for determining parasite persistence. However understanding the key parasite characteristics and host pathways driving these patterns is important, as resource provision can impact infectious disease dynamics in opposite directions. For instance, population increases in vector-borne systems could lead to increments in parasite prevalence. Hence it is crucial to recognise system-specific variations, such as specific host behaviours at the individual level, parasite interactions with the host immune system, and how natural resource availability shapes host demographics, in order to fully understand and predict epidemiological responses to provisioning.

## Data Availability Statement

Data available from https://github.com/dieraz/prov-theo.git

## Acknowledgements

We would like to thank those involved in the fieldwork that generated the data used in this study, and the landowners for permission to carry out the work on their land. The work was funded by NERC Grants NE/G007349/1 and NE/G006830/1, NE/I024038/1, NE/I026367/1 and NE/R011397/1, awarded to AF and ABP. ABP was funded by a University of Edinburgh Chancellors Fellowship and a Wellcome Trust Grant (095831).

