## Supplemental material for "The impact of resource provisioning on the epidemiological responses of different parasites"

**Supplementary material**

*Model frameworks*

Each model had, at its heart, the same host demographic model:

$\frac{dN}{dt}=bN\frac{(K-N)}{K}-\mu N$	Eq S1

where$N$ represents the total host population, $K$ the carrying capacity, $b$the birth rate and, $\mu$ the mortality rate; in what follows we assumed infections do not increase host mortality. For our analyses, we used this demographic model as a template, and developed five compartmental models, each reflecting different parasites types (inspired by our field system; described below), to study the effects of host resource provisioning on the dynamics of each parasite. Each model incorporated parasite transmission parameters reflecting contact behaviour ($\alpha:$ contact rate) and immune defence ($\delta:$ host susceptibility), that were allowed to vary with provisioning (see main text).

Our first microparasite transmission model is the **SIR model**, in which $S$, $I$, and $R$ represent the susceptible, infected and recovered host populations. Susceptible hosts become infected when having an encounter with infected hosts, according to the term $\alpha\delta SI$, where $\alpha$ is the per capita encounter rate between susceptible and infected hosts, and $\delta$ is host susceptibility or the probability that the encounter results in infection. Infected individuals recover to life-long immune hosts at rate $\gamma$. The SIR model is therefore represented by a two-equation system (where the dynamics of the *R* class are implicit, as *N* = *S+I+R*), as follows:

$\frac{dS}{dt}= b N\frac{(K-N)}{K}-\alpha\delta SI+\gamma I-\mu S$	Eq S2

$\frac{dI}{dt}= \alpha\delta SI-\gamma I-\mu I$	Eq S3

For the **microparasite environmentally-transmitted SIS model**, as motivated by an *Eimeria* (gastrointestinal coccidia) parasite (Table 1, main text), hosts get infected when ingesting eggs in the environment ($E$), according to the term $\alpha\delta SE$, where $\alpha$ is the per capita encounter rate between hosts and eggs, and $\delta$ is host susceptibility or the probability that the encounter results in infection. Each infected individual sheds eggs at rate $\tau$ and recovers (back to being a fully susceptible host) at rate $\gamma$. Eggs in the environment could become inactive and lose the ability to infect susceptible hosts, thus $\phi$represents egg decay rate in the environment. The microparasite environmentally-transmitted (SIS) model is therefore represented by a three-equation system as follows:

$\frac{dS}{dt}= b N\frac{(K-N)}{K}-\alpha\delta SE+\gamma I-\mu S$	Eq S4

$\frac{dI}{dt}= \alpha\delta SE-\gamma I-\mu I$	Eq S5

$\frac{dE}{dt}= \tau I-\phi E$	Eq S6

For the **sex-biased microparasite SAL model**, as motivated by a general herpesvirus system (Erazo, Pedersen, Gallagher, & Fenton, 2021), susceptible individuals ($S$) become infected and enter the active class ($A$) through contact with active-infected individuals (latent-infected individuals are assumed not to transmit infections) at transmission rate $\alpha\delta$, where $\alpha$ is the per capita encounter rate between susceptible and acute hosts, and $\delta$ is host susceptibility or the probability that the encounter results in infection. Active-infected individuals then move to the latent class ($L$) at transition rate $\varepsilon$ (hence active infections last on average 1/$\varepsilon$ weeks), and vice versa at rate $\eta$ (hence latent infections last on average 1/$\eta$ weeks). We assumed individuals do not recover from infections.

The model considered explicit female and male classes and density-dependent transmission for all possible transmission routes ($\beta_{1}-\beta_{4}$). Therefore, the model consisted of six classes (susceptible, active-infected and latent-infected, for both females and males), and all transmission terms had the form ${\alpha_{n}\delta}_{n}S_{i}A_{j}$ where $i$ and $j$ represent sex, either female or male, and ${\alpha_{n}\delta}_{n}$ is the transmission term between $i$ and $j$. The sex-biased microparasite model is therefore represented by a six-equation system as follows:

$$\frac{dS_{m}}{dt}= b \frac{N}{2}\frac{(K-N)}{K}-\alpha_{1}\delta_{1}S_{m}A_{m}{-\alpha}_{2}\delta_{2}S_{m}A_{f}-\mu S_{m}$$

 Eq S7

$\frac{dA_{m}}{dt}=\alpha_{1}\delta_{1}S_{m}A_{m}{+\alpha}_{2}\delta_{2}S_{m}A_{f}-\mu A_{m}-\varepsilon A_{m}+\eta L_{m}$	Eq S8

$\frac{dL_{m}}{dt}=\varepsilon A_{m}-\eta L_{m}-\mu L_{m}$	Eq S9

$\frac{dS_{f}}{dt}= b \frac{N}{2}\frac{(K-N)}{K}-\alpha_{3}\delta_{3}S_{f}A_{m}{-\alpha}_{4}\delta_{4}S_{f}A_{f}-\mu S_{f}$	Eq S10

$\frac{dA_{f}}{dt}=\alpha_{3}\delta_{3}S_{f}A_{m}{+\alpha}_{4}\delta_{4}S_{f}A_{f}-\mu A_{f}-\varepsilon A_{f}+\eta L_{f}$	Eq S11

$\frac{dL_{f}}{dt}=\varepsilon A_{f}-\eta L_{f}-\mu L_{f}$	Eq S12

The **vector-borne SIS system** considers vector ($S_{v}$, $I_{v})$ and host populations ($S_{h}$, $I_{h})$, which are divided by their infection status. The vector population increases at rate $b_{v} N_{v}\frac{(K_{v}-N_{v})}{K_{v}}$ where $b_{v}$, $N_{v}$ and $K_{v}$ represent vector birth rate, total vector population and vector carrying capacity, respectively. Susceptible vectors ($S_{v})$ become infected when having a successful encounter with infected hosts $(I_{h})$, expressed by the $\beta S_{v}I_{h}$ term where $\beta$ is the host-to-vector transmission rate. Vectors die at a $\mu_{v}$ rate. Susceptible hosts ($S_{h}$) become infected through contact with infected vectors ($I_{v})$ at vector-to-host transmission rate $\alpha\delta$, where $\alpha$ is the per capita encounter rate between susceptible hosts and infected vectors, and $\delta$ is host susceptibility or the probability that the encounter results in infection. Infected hosts $(I_{h})$ recover at a $\gamma$ rate. The SIS vector-borne model is therefore represented by a four-equation system as follows:

$\frac{dS_{v}}{dt}= b_{v} N_{v}\frac{(K_{v}-N_{v})}{K_{v}}-\beta S_{v}I_{h}-{\mu_{v}S}_{v}$	Eq S13

$\frac{dI_{v}}{dt}= \beta S_{v}I_{h}-{\mu_{v}I}_{v}$	Eq S14

$\frac{dS_{h}}{dt}= b N_{h}\frac{(K_{h}-N_{h})}{K_{h}}-\alpha\delta S_{h}I_{v}-\mu S_{h}+\gamma I_{h}$	Eq S15

$\frac{dI_{h}}{dt}= \alpha\delta S_{h}I_{v}-\mu S_{h}-\gamma I_{h}$	Eq S16

In contrast with the microparasite models, the **macroparasite model** keeps track explicitly of the number of macroparasites $(P$) within hosts $(H$), which reproduce and release larvae $(L$) into the environment. Adult parasites $(P$) are assumed to be acquired when hosts $(H$) have an encounter with the parasite free-living stage in the environment. $P$ decreases by two factors: host natural mortality ($\mu$) and host parasite-induced mortality ($\nu P$) dependent on the mean burden of worm infection. For macroparasites parasite aggregation or crowding is incorporated via the term $\frac{P^{2}\nu(k+1)}{Hk}$ where $k$ represents the aggregation coefficient (exponent of the negative binomial distribution) (Anderson & May, 1978). The environmental pool of larvae $(L$) increases when the adult parasites $(P$) release larvae at $\tau$ rate and decreases due to death at rate $\phi$. The macroparasite model is therefore represented by a three-equation system as follows:

$\frac{dH}{dt}= b H\frac{(K-H)}{K}-\mu H-\nu P$	Eq S17

$\frac{dP}{dt}= \alpha\delta LH-\left( \mu+\nu\right)P-\frac{P^{2}\nu\left( k+1 \right)}{Hk}$	Eq S18

$\frac{dL}{dt}= \tau P-\phi L$	Eq S19

*Field data collection for model parameterisation*

Data from individuals was generated longitudinally from wood mice (*Apodemus sylvaticus*) collected between June 2009 and December 2012 in four grids located in Haddon Wood, Cheshire, UK as described in (Knowles, Fenton, & Pedersen, 2012; Sweeny, Albery, Venkatesan, Fenton, & Pedersen, 2021). Two Sherman traps baited with grain and bedding were placed every 10 m in each 70 x 70 m squared grid. Trapping was conducted for 3 consecutive nights every 3 weeks during four field seasons: June-December in 2009 and 2012, and May-December in 2010 and 2011. All trapped individuals were tagged for recognition in subsequent recaptures. From all wood mice, morphometric measures were recorded, including sex and age.

The main parasite species in this system have been well characterised (Knowles et al., 2012, 2013; Sweeny et al., 2021; Withenshaw, Devevey, Pedersen, & Fenton, 2016), and are summarised in Table 1. Methods of identification of each parasite are described in the above references, but briefly, the GI parasites *H. polygyrus* and *E. hungaryensis* were identified from eggs or oocysts in faecal samples collected from the traps using the salt flotation technique; infection levels were quantified as faecal egg or oocyst counts (Knowles et al., 2013). The presence of microparasites were determined from blood samples taken from the tail at first capture within each month. Wood mouse Herpes Virus and Cowpox infection was detected using a serological assay that detects antibodies in mouse serum using IFA (Knowles et al., 2012). Bacteria of the genus *Bartonella*, and *Trypanosoma grosi* were screened and identified using PCR-based diagnostics (Withenshaw et al., 2016). All parasitological data was aggregated weekly. Mouse population size per week was defined as the total number of mice collected in the four grids during that period.

*Model parameterization*

Demographic parameters ($\sigma_{d}=\{K, \mu, b\})$ and parasite-related parameters ($\sigma_{p}$, which varied depending on the model type being fitted; see below) were estimated using adaptive Monte Carlo Markov Chain Metropolis-Hastings (MCMC-MH) (Camacho & Funk, 2019), assuming uniform priors, through fitting to data on wood mouse population abundances and infection seroprevalences or prevalence for microparasites and faecal/oocyst eggs counts for macroparasites. We ignored the first year (52 weeks) of predicted transient dynamics of the simulation as burn-in time, and fitted the models over the subsequent 4 years of data.

Model fitting was carried out in two stages. First, demographic parameters and carrying capacity were estimated by fitting the simulated total number of weekly wood mice (*l_mice_*; $N$ from Eqn S1) to the observed number of mice captured per week (*y_week_*). The weekly number of wood mice captured was assumed to follow a Poisson distribution. The log-likelihood of the data for the mice population dynamics model was given by:

$$l_{mice}(\sigma_{d})=l_{mice}\left( data | \sigma_{d} \right)=\sum_{week} l_{mice-week}(y_{week}|\sigma_{d})$$

Through this we generated posterior distributions for the carrying capacity, birth and death rate.

Next, we sought to estimate the different parasite-related parameters in the various parasite transmission models. Models were parameterized from data on wood mice infected with five parasites: *Eimeria hungaryensis,* Wood Mouse Herpes Virus*, Bartonella* spp., *Trypanosoma grosi* and *Heligmosomoides polygyrus.* Note that the transmission rate for all parasites is the product of contact rate ($\alpha)$ and susceptibility ($\delta)$, thus for model fitting each of these products were considered as one single parameter for each parasite. For the subsequent provisioning analysis these transmission term parameters were assumed individually as defined in the model construction, because resource provisioning may induce different responses in direction and magnitude for contact rate versus susceptibility.

Disease-related parameters estimated for each system were: *E. hungaryensis*:$\sigma_{p}=\{\alpha\delta,\gamma\}$, Herpesvirus:$\sigma_{p}=\{\alpha_{1}\delta_{1},\alpha_{2}\delta_{2},\alpha_{3}\delta_{3},\alpha_{4}\delta_{4}, \varepsilon,\eta\}$, *Bartonella* spp. and *T. grosi:*$\sigma_{p}=\{\beta,\alpha\delta,\gamma\}$, $=\{\alpha\delta,\gamma\}$, and *H. polygyrus:*$\sigma_{p}=\{\alpha\delta,\nu\}$. For the microparasites, the weekly simulated number of infected mice was fitted to the observed number of mice infected by that parasite per week. The observed number of infected mice were assumed to follow a Poisson distribution. The log-likelihood of the data is given by:

$$l\left( data | \sigma_{p} \right)=\sum_{week} l_{prev-week}\left( y_{week} | \sigma_{p} \right)$$

For the macroparasite, the weekly simulated total *H. polygyrus* larvae released by mice was fitted to the total larval counts in faecal samples per week. The number of larvae was assumed to follow a negative binomial distribution. The log-likelihood of the data is given by:

$$l\left( data | \sigma_{p} \right)=\sum_{week} l_{larvae-week}(y_{week}|\sigma_{p})$$

Three MCMC-MH chains of 10,000 iterations per model were run using the default parameter standard deviation (parameter value divided by 10). Then, for each chain, using the first chain output (standard deviation and $\bar{\theta}$for the last 9,000 iterations) as input, we ran a second chain of 100,000 iterations. For the second chain, the first 5,000 iterations were discarded (burn-in), and we eliminated every 10 samples per sample to avoid auto-correlation (thinning). The Gelman-Rubin diagnostic was used to assess MCMC convergence by analysing the difference between chains. Through this we first generated posterior distributions for the carrying capacity, birth and mortality rate. Using this demographic parameter estimates, we then generated posterior distributions for disease-related parameters for each parasite system individually.

Since cowpox infections were so rare in our system (<5%), we estimated its recovery rate and prevalence in wood mice from Telfer et al. (2002); they reported prevalence in wood mice ranging $0\%-17\%$, we assumed infection in wood mice is on average $0.085$. For obtaining this average wood mice prevalence in the SIR system, we found that transmission rate was $0.013\frac{1}{ind \cdot week}$.

**Parameter estimation results**

Host birth rate was estimated to be $1.24 [1.23-1.25]$ $\frac{ind}{ind \cdot week}$and mortality rate was $\frac{1}{13.75 [13.57-13.94] weeks}$. The estimated carrying capacity was $42.12 [42.05-42.19]$ mice per grid. All estimated disease-related parameters are shown in Table S2.

| Parasite | Transmission rate | Recovery rate  $\boldsymbol{(\gamma)}$ | Disease-Induced mortality $\boldsymbol{(\nu)}$ | Acute to latent transition rate $\boldsymbol{(\varepsilon)}$ | Latent to acute transition rate $\boldsymbol{(\eta)}$ |
| --- | --- | --- | --- | --- | --- |
| Cowpox virus  (SIR) | $\alpha\delta=0.013$ | $0.25$  (Telfer et al., 2002) |  |  |  |
| *Eimeria hungaryensis*  (SIS) | $\alpha\delta=9.59x{10}^{-7}$  $[9.58-9.61]$ | $0.1131$  $[0.1128-0.1134]$ | - |  |  |
| Herpesvirus  (SAL) | Male to male:  $\alpha_{1}\delta_{1}=0.0255$  $[0.0253-0.0256]$  Female to male:  $\alpha_{2}\delta_{2}=0.0424$  $\left[ 0.0421-0.0428 \right]$  Male to female:  $\alpha_{3}\delta_{3}=0.0148$  $\left[ 0.0147-0.0149 \right]$  Female to female:  $\alpha_{4}\delta_{4}=0.0156$  $\left[ 0.0154-0.0157 \right]$ |  |  | $0.815$  $[0.813-0.817$ | $0.0132$  $[0.0131-0.0133$ |
| *Bartonella* spp.  (vector-borne) | Host to vector:  $\beta=0.506$  $[0.503-0.509]$  Vector to host:  $\alpha\delta=8.15x{10}^{-3}$  $[8.11-8.18]$ | $0.643$  $[0.639-0.6459]$ |  |  |  |
| *Trypanosoma grosi*  (vector-borne) | Host to vector:  $\beta=0.547$  $[0.544-0.551]$  Vector to host:  $\alpha\delta=1.056x{10}^{-3}$  $[1.052-1.061]$ | $0.644$  $[0.641-0.647]$ |  |  |  |
| *Heligmosomoides polygyrus*  (macroparasite) | $\alpha\delta=2.05x{10}^{-4}$  $[2.04-2.06]$ | - | $0.765$  $[0.763-0.767]$ |  |  |

**Table S2. Parasite-related parameters. Estimated values show median and 95% credible intervals from the posterior distributions of each parameter.**

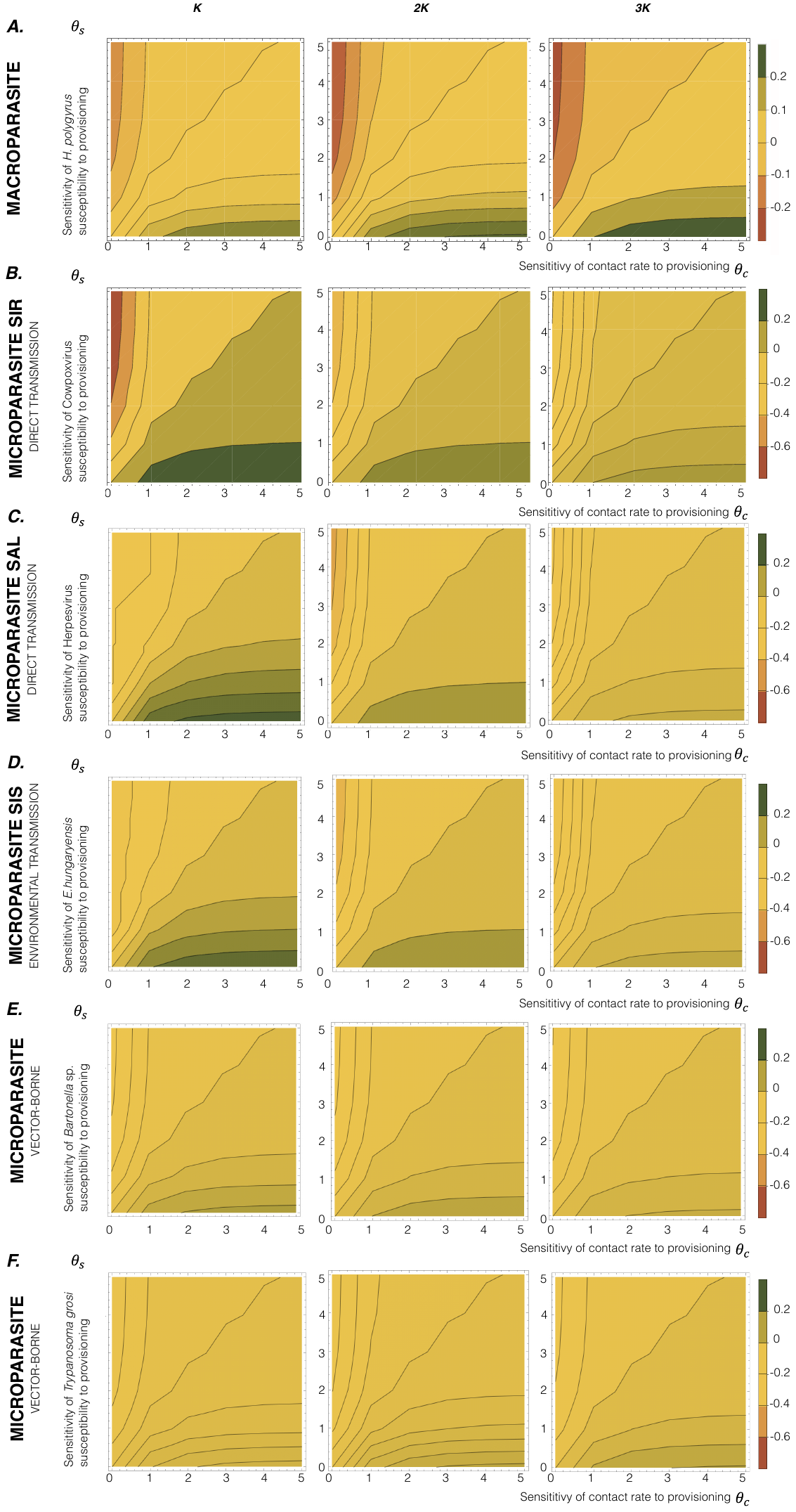

**Figure S1. Parasite outcome changes induced by provisioning.** A row shows macroparasite mean burden change and B row to F row represent normalized microparasites’ prevalence changes (the difference in equilibrium prevalence between provisioned ($\rho=1$) and un-provisioned scenarios ($\rho=0$), adjusted to the same scale across all parasites). Columns illustrate different carrying capacities, varying from $K$ to $3K$, representing increasing levels of host demographic response to provisioning. For all figures, *X-*axis represents the sensitivity of contact rate to provisioning $(\theta_{c})$ and *y-*axis is the sensitivity of host susceptibility to provisioning $\left( \theta_{s} \right).$
